# Rho (ρ) Analysis to Dissect RNA Folding and Assembly Pathways

**DOI:** 10.64898/2026.07.18.739347

**Authors:** Brant Gracia, Sarah E. Nielson, Daniel Herschlag, Rick Russell

## Abstract

The modular structure and energetics of RNA simplifies its folding. Leveraging this modularity, we introduce Rho (ρ) analysis to systematically dissect RNA conformational pathways. ρ analysis uses isolated RNA secondary or tertiary contacts as external standards to provide insights not possible via the “internal” comparisons of traditional ϕ analysis. Equivalent effects of a mutation on the folding rate constant of the RNA of interest and the thermodynamic stability of the isolated contact indicate that the mutated interaction is fully formed prior to the rate-limiting transition state; the absence of a kinetic effect indicates that the interaction is formed after this transition state. Comparisons with properties of the isolated contact provide additional insights about conformational pathways. We demonstrate ρ analysis by dissecting *Tetrahymena* group I intron folding pathways, using a split intron in which the P5abc subdomain assembles with the intron core through three tertiary contacts. We uncover multiple folding pathways and modulation in pathway flux that are readily understood from the energetic properties of the constituent RNA motifs. Extending these concepts to RNA-guided DNA recognition by CRISPR-Cas12a, crRNA–DNA mismatches give substantial ϕ values across much of the target, indicating a late transition state in binding. Thermodynamic penalties from mismatches support modular base-pairing energetics and define an upper bound on DNA target specificity. Our results establish ρ analysis as a general framework to probe RNA conformational pathways and function. It is straightforward to implement and can be readily applied *in vitro* and in cells.

## INTRODUCTION

Structured RNAs play essential roles throughout biology. To form their active states and to function, RNAs traverse free energy landscapes that include multiple on-pathway and off-pathway intermediates, with barriers that produce transitions on time scales that span more than six orders of magnitude and routinely produce long-lived misfolded states (1–3). Superficially, this behavior seems more complex than that for protein folding, which has been intensively studied for decades but where a predictive understanding of folding pathways, rates and stabilities is still lacking (4, 5). However, RNA folding studies have revealed simplifications in its structure and energetics, in particular in the modularity and sparseness of RNA tertiary contacts that in turn has led to a general, predictive model for RNA folding kinetics and thermodynamics ((6–8); reviewed in (8)).

According to the RNA Reconstitution Model, preformed helices and junctions (so-called helix-junction-helix elements) undergo a biased probabilistic search to allow tertiary contact partners to find one another and form a contact that stabilizes the folded state. For RNAs with multiple tertiary contacts, this process is repeated for the formation of each contact, with different pathways specifying different formation orders and with any of the steps potentially highest in energy and rate-limiting within a pathway.^‡^ The Reconstitution Model and the simplifying RNA features that support it have allowed us to develop Rho (ρ) analysis herein to dissect these folding pathways and determine the order of formation of RNA contacts, rate-limiting steps, and additional information about these pathways.

ρ analysis builds on rate-equilibrium free energy relationships (9), which were applied to protein folding by Fersht as Phi (ϕ) analysis (10). For ϕ analysis, the destabilizing effect of a mutation on the rate-limiting transition state, relative to its destabilization of the folded state, provides a measure of the extent to which the native contact is present in the transition state. ϕ analysis has been powerful for dissecting protein folding, but the extensive, complex networks of cooperative interactions that are inherent to globular protein folds change throughout folding by the progressive amalgamation of interaction networks in later folding states, complicating the energetic relationships underlying ϕ analysis (11–13).

The energetic modularity observed for RNA suggests that complicating network effects will be rare for RNA, simplifying ϕ analysis for RNA. Further, if energetic modularity holds for RNA, we should be able to use the kinetic and thermodynamic effects of mutations on *isolated motifs as an external standard*. Comparison of the effects of the same mutations on folding of a complex RNA to their effects on a simpler RNA containing only that tertiary motif should reveal when along the folding landscape that interaction is made (i.e., before or after the transition state). Additional information can be obtained about what step is rate limiting, what folding pathway is favored and whether multiple pathways are followed, and whether the tertiary motif is perturbed in the final folded state, as described in the “Approach” section below and with our results. RNAs also undergo conformational transitions in carrying out their functions, so that ρ analysis may also be used to dissect and understand reaction pathways and their underlying energetics.

To develop and test ρ analysis, we chose two systems: one involving RNA tertiary structure formation and the other involving secondary structure formation. Both systems have been extensively characterized by structural and biophysical studies, providing well-established structural frameworks for ρ analysis. We first applied ρ analysis to tertiary assembly of the *Tetrahymena* group I intron ribozyme by comparing the energetic effects of mutations during ribozyme assembly with analogous measurements of an isolated tertiary contact as an external energetic reference (6, 7, 14). This analysis revealed multiple folding pathways and showed that mutations can redirect pathway flux by altering the energetic accessibility of individual tertiary contacts. We next applied ρ analysis to secondary structure formation by analyzing RNA-guided DNA recognition by CRISPR- Cas12a through formation of RNA-DNA base pairs (an R-loop). In this system, the energetic effects of mismatches support modular base-pairing energetics, reveal a late transition state for R-loop formation, and define an upper limit on DNA target specificity. Together, these results show how the intrinsic energetics of modular RNA structural elements and the placement of transition states define RNA conformational landscapes and shape RNA folding and molecular recognition. Further, they establish ρ analysis as a general framework for quantitatively dissecting those landscapes.

### APPROACH: ρ Analysis to Dissect RNA Folding and Assembly

The ϕ analysis formalism was developed for protein folding and assembly (10), and it has been applied (on a more limited basis) to RNA folding (15, 16). Immediately below we formally define these values and the ρ values that are introduced herein. We then describe the information that can be obtained from these parameters.

ϕ (Eq. 1): This term is the standard value defined by Fersht to describe the effect of a mutation on the folding rate constant relative to that mutation’s effect on thermodynamic stability of the protein or RNA of interest. In the simplest scenario, ϕ = 1 or 0 if the mutated residue’s interactions are made before or after (respectively) the rate-limiting transition state for folding. Intermediate values are more difficult to interpret, as they can arise from multiple folding pathways or from local interactions that are formed but give different energetic effects in the partially-folded transition state than in the fully-folded macromolecule (11–13).

ϕ, like all of the comparisons, is made in terms of free energies, and, for clarity, we refer to these free energy parameters with a “≠” superscript for rate effects and with an “eq” superscript for equilibrium effects in Eq. 1; the subscript “ROI” denotes “RNA of interest” and is used to distinguish measurements with the ROI from measurements with external comparisons (subscripted “ref” for “reference”), which are used in ρ analysis (Eqs. 2 & 3 below).

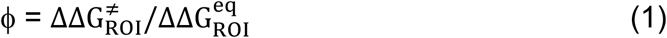

ρ_ref_ (Eq. 2): This term is the same as a standard ϕ value but for the specific case of an RNA tertiary contact in isolation (Eq. 2; Fig. 1A). These “reference” values can be compared to ϕ values for the RNA of interest (ROI) to assess whether the overall folding rate of the ROI is limited by formation of this tertiary contact. In addition, the denominator in the expression for ρ_ref_ is used in obtaining ρ_ext_ (Eq. 3 below).

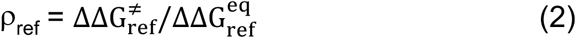

**Figure 1.**
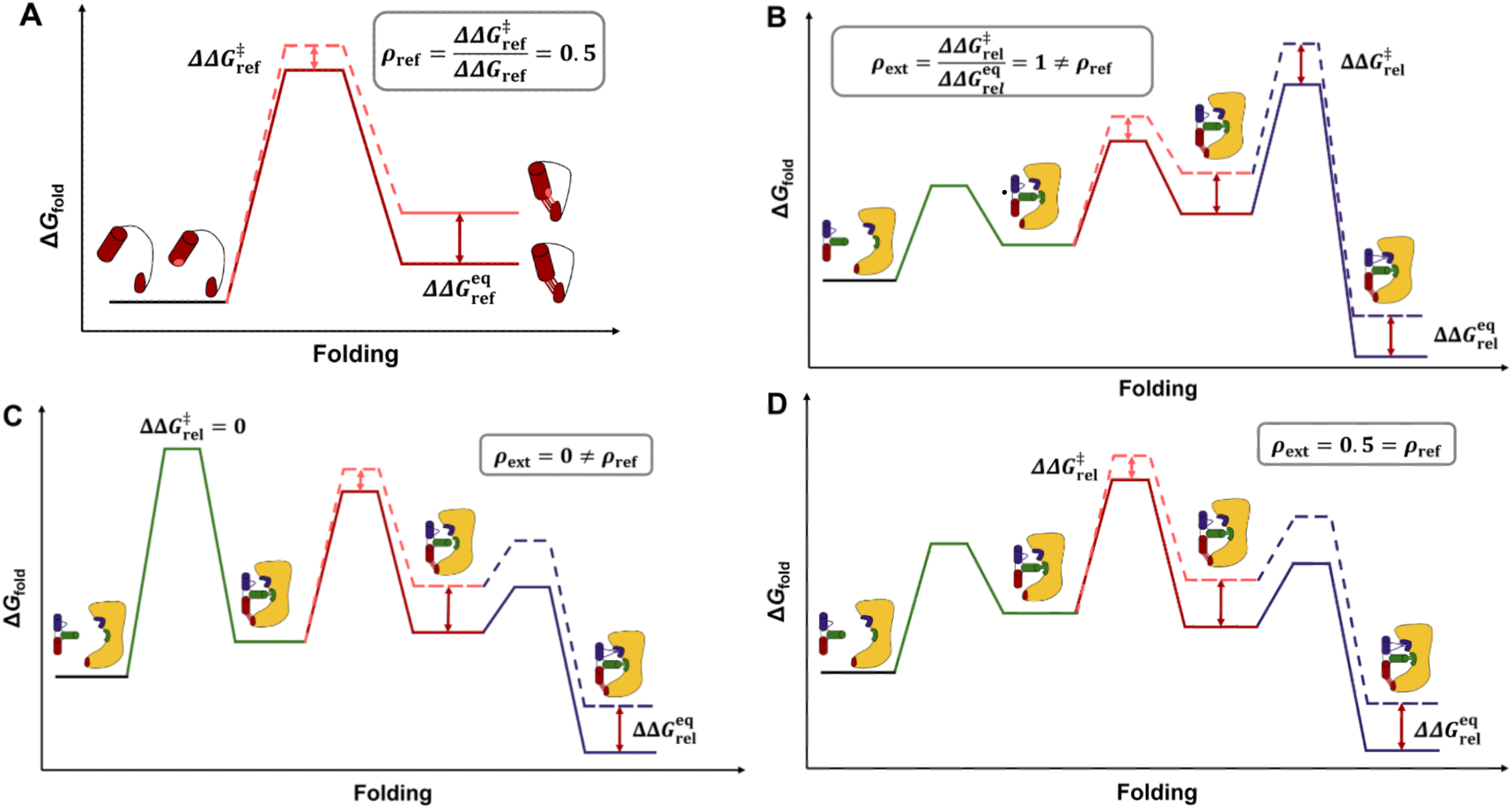
Principles of ρ analysis. **A.** In ρ analysis, the kinetic and thermodynamic effects of mutations on folding of the isolated tertiary contact are used to generate ρ_ref_. The ρ_ref_ value is shown as equal to 0.5 in this example, as the free energy effect of the mutation (depicted with lighter shaded region) on the rate constant is half of the effect on the equilibrium constant. This contact and mutation are used in the examples in panels **B–D**. **B–D.** Hypothetical folding of a complex RNA. The assembly of the *Tetrahymena* group I intron studied herein is schematically shown, with the yellow representing E^ΔP5abc^ and the three colored, connected helices representing P5abc (see Fig. 2 & text for definitions). For purposes of illustration, Panels **B–D** show a common pathway (order of tertiary contact formation), with each panel depicting a different rate-limiting step. The ρ value for the contact from Panel A (shown in red) is given for each free energy profile. **B**: The rate- limiting step is the *last* step, after formation of the red contact, therefore giving ρ_ext_ = 1. **C**: The rate-limiting step is the *first* step, before formation of the red contact, therefore giving ρ_ext_ = 0. **D**: The rate-limiting step is the *middle* step, corresponding to formation of the red contact, therefore giving ρ_ext_ = ρ_ref_ = 0.5.

ρ_ext_ (Eq. 3): ρ_ext_ compares the effect of a mutation on the folding or assembly rate for the RNA of interest (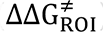) to the equilibrium effect of that same mutation for the contact in isolation (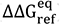). For complex RNAs and RNA•protein assemblies, two-state folding often does not hold and overall thermodynamic effects (which are needed for classical ϕ analysis) can be difficult or impossible to determine; ρ_ext_ has the advantage of eliminating the need to measure the equilibrium folding effect of each mutant in the ROI (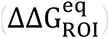). Instead, it uses a universal reference value (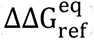), a value that must be measured for tertiary contacts but can be calculated from nearest-neighbor rules for base pairs. Using these reference values allows ρ analysis to provide information analogous to—but even more powerful than—traditional ϕ analysis, as described by the examples in Fig. 1 below and as illustrated by our experimental results.

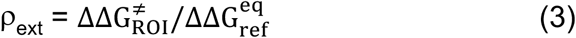

Fig. 1B–D depicts an RNA folding process involving the formation of three tertiary interactions. The three panels show different folding landscapes that give different values of ρ_ext_ and thus can be distinguished by measuring ρ_ext_. Fig. 1A shows the folding of an RNA with only one of the tertiary interactions (in isolation) that is used as the reference to obtain ρ_ref_ and is then used in calculating the ρ_ext_ value. Measuring ρ_ref_ ensures that the mutation to be studied affects folding, and determining the size of the effect allows the quantitative comparisons needed to infer the nature of the folding landscape.

If the interrogated tertiary contact forms *prior to* the rate-limiting transition state and fully mimics the interaction in the final state, mutations that weaken that tertiary contact will slow assembly *to the same extent* that they (thermodynamically) destabilize the contact in isolation (ρ_ext_); they will not affect the dissociation rate constant. These effects give ρ_ext_ = 1 (Fig. 1B). Conversely, weakening a contact that forms after the rate-limiting transition state will not affect the assembly rate constant so that ρ_ext_ = 0 (Fig. 1C). (Instead this mutation will increase the reverse process—disassembly or unfolding—by the amount that is equal to the destabilization of the final structure so that ρ_ext_ for unfolding equals 1; the sum of ρ_ext,folding_ + ρ_ext, unfolding_ equals 1, by definition.) Finally, if the tertiary contact is *formed in the rate-limiting transition state*, and if the contact forms in the same way in the complex RNA as it does when isolated, then ρ_ext_ will equal ρ_ref_ (Fig. 1D). The measurements with the isolated tertiary interaction (ρ_ref_) allow intermediate ρ_ext_ values to be interpreted and provide information about the rate-limiting transition state.

In practice, having multiple mutations for each tertiary interaction strengthens the conclusions that can be drawn, and additional information about folding pathways can be obtained by making the same set of mutations in different backgrounds—e.g., with other tertiary interactions altered or ablated. Below, we apply ρ analysis to a model RNA folding and assembly process and then to an RNA-guided DNA recognition process in a CRISPR-Cas endonuclease. Our ρ analysis provides incisive information about pathways and specificity that would be difficult or impossible to obtain without this new approach.

## RESULTS AND DISCUSSION

To test ρ analysis and dissect a complex RNA tertiary assembly process, we studied the folding of the *Tetrahymena* group I intron, using a split intron in which the P5abc subdomain assembles with the ribozyme core (E^ΔP5abc^) through three modular tertiary contacts (17, 18): a tetraloop/tetraloop receptor (TL/TLR) interaction, a metal core/receptor (MC/MCR) interaction, and a kissing loop interaction, termed P14 (Fig. 2A, colored arrows). The TL/TLR and MC/MCR interactions form within the P4-P6 domain and have been the focus of extensive previous structural and biophysical work [(6, 19) and references therein], and P14 forms between the L5c loop within P5abc and the L2 loop within E^ΔP5abc^.

**Figure 2.**
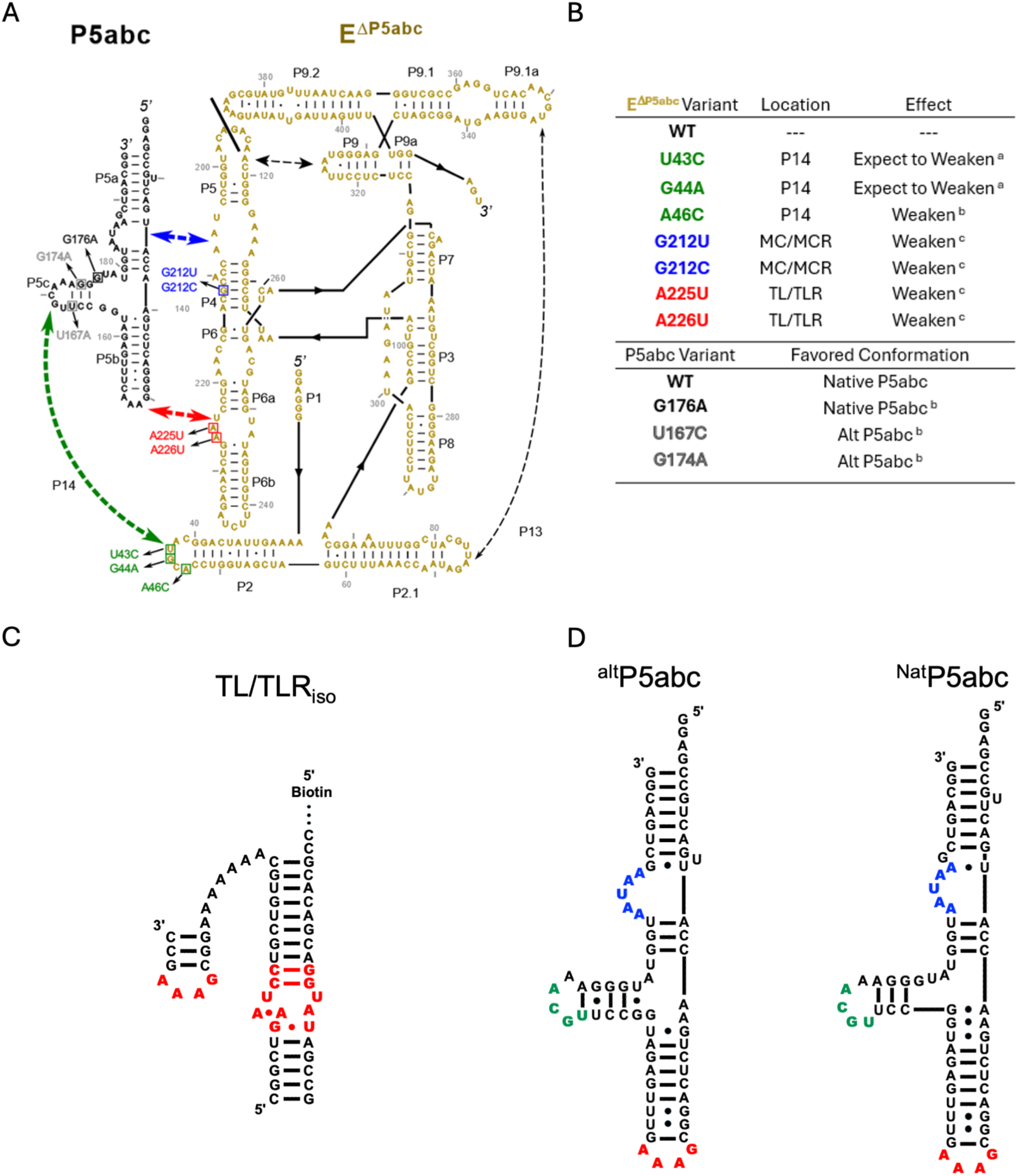
P5abc and E^ΔP5abc^ RNAs: a two-component version of the *Tetrahymena* group I ribozyme. **A.** RNA secondary structures and tertiary contacts in the assembled E^ΔP5abc^•P5abc complex. P5abc RNA is shown on the left in black and E^ΔP5abc^ on the right in orange. The three tertiary contacts formed between the two RNAs are indicated and color coded, with these colors used in subsequent figures. The P5abc and E^ΔP5abc^ mutations used herein are indicated. **B.** P5abc and E^ΔP5abc^ variants used herein are listed using the same color scheme as in panel **A** for ease of reference along with their expected effects. (^a,^ ref. (46); ^b,^ ref. (20); ^c,^ ref. (6)) **C.** Schematic of the TL/TLR^iso^ construct (6, 47). **D.** Secondary structures of misfolded (^alt^P5abc, left) and native (^Nat^P5abc, right) P5abc.

Using an electrophoretic mobility shift assay (EMSA), we measured the kinetics of P5abc binding and dissociation with the wild-type E^ΔP5abc^ and two TLR mutants that were previously characterized in an isolated TL/TLR system (A225U and A226U; Fig. 2A,B,C) (6). The measurements in the isolated system provide kinetic and equilibrium binding constants and the reference thermodynamic values, 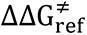 and 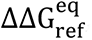, for ρ analysis (Eq. 2).

We used ρ analysis to determine the assembly pathway for WT and mutant P5abc RNAs. U167C and G174A P5abc favor an alternative structure, termed ^alt^P5abc, and slow assembly with E^ΔP5abc^ (20), and we wanted to know the origin of this effect and whether it arose from slowing a common folding pathway or enforcing an alternative, slower pathway. ^alt^P5abc includes a shift in the register of the short P5c helix and loss of native structure in the three-helix junction within P5abc (Fig. 2D (20–26)). We first present the results for these P5abc mutants as they provide a straightforward deomonstration of ρ analysis.

### ρ analysis reveals rate-limiting formation of the TL/TLR contact

Using standard kinetics methods and EMSA to separate P5abc free and bound to E^ΔP5abc^, we measured binding and dissociation kinetics for each of the two mutants that stabilize ^alt^P5abc (U167C and G174A). In these P5abc mutant backgrounds, the TLR mutations slowed association (10-18-fold; Fig. 3A; Fig. S1), accelerated dissociation (5- 13-fold; Fig. 3B; Fig. S2), and destabilized the complex overall (40-120-fold, corresponding to 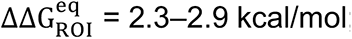; Fig. 3C).

**Figure 3.**
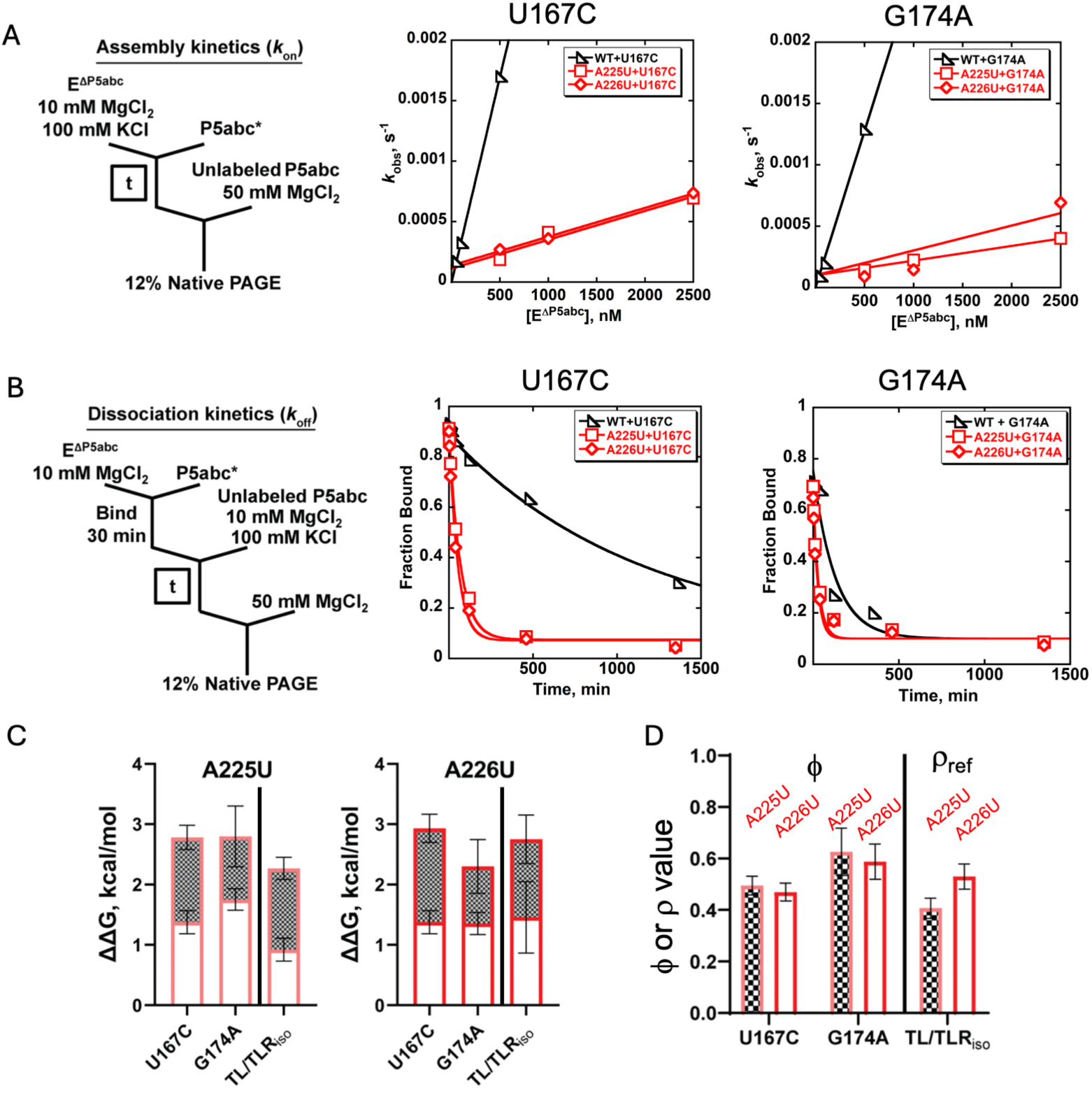
ρ analysis with stabilized ^alt^P5abc variants reveals rate-limiting TL/TLR formation in the assembly of P5abc and E^ΔP5abc^. **A.** Assembly kinetics. Left, the experimental scheme for measuring assembly kinetics. Center and right, dependences of observed rate constants for P5abc binding (U167C and G174A, respectively) on the concentration of E^ΔP5abc^, for WT and the indicated E^ΔP5abc^ mutants. **B.** Dissociation kinetics. Left, the experimental scheme for measuring dissociation kinetics. Center and right, dissociation time courses for P5abc (U167C and G174A, respectively) from E^ΔP5abc^ with WT and the indicated E^ΔP5abc^ mutants. **C.** Energetic effects of the TL/TLR mutations (A225U and A226U) on P5abc/E^ΔP5abc^ assembly (white bars) and disassembly (hatched bars) kinetics. The bars are stacked to illustrate the overall (thermodynamic) energetic effects of the TL/TLR mutations (relative to WT). The effects of the same mutations on the isolated TL/TLR contact are shown at the right in each graph (6). **D.** ϕ values for the A225U and A226U TL/TLR mutations (hatched and solid bars, respectively) in the context of U167C and G174A P5abc association with E^ΔP5abc^ match the ρ values for these mutations in the isolated TL/TLR (ρ_ref_; (6)).

The total amount of destabilization was the same, within error, as the corresponding destabilization of the TL/TLR contact in isolation for each mutation (Fig. 3C). This equivalence supports modular energetics of the TL/TLR contact, as predicted by the RNA reconstitution model (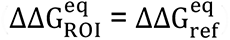). Strikingly, the effects of the TLR mutations on the binding and dissociation kinetics were also the same as those for TL/TLR formation in isolation (Fig. 3C; 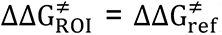). Observing the same values (i.e., the same kinetic and thermodynamic effects) for the complex assembly of ^alt^P5abc•E^ΔP5abc^ for the isolated TL/TLR construct strongly suggests that TL/TLR formation is rate-limiting for assembly and unperturbed in the final complex (see Fig. 1D). In formal terms, these observations are equivalent to showing that ϕ equals ρ_ref_ (Eq. 2 & 3).

The U167C and G174A mutations give ϕ values of 0.5 and 0.6, respectively (Fig. 3D). In traditional analysis, values between 0 and 1 are difficult to interpret, as they can represent partially rate-limiting TL/TLR formation, multiple folding pathways, or formation prior to the rate-limiting step that are followed by additional changes that strengthen the TL/TLR. However, the external standards in ρ analysis (Eq. 2 & 3) allow the intermediate ϕ values to be interepreted. Specifically, the observed ϕ values match those for the isolated reference for TL/TLR formation (ϕ = ρ_ref_), providing strong evidence that TL/TLR formation is the rate-limiting step in ^alt^P5abc•E^ΔP5abc^ assembly and forms similarly to its formation in isolation, as noted above. TL/TLR contacts, which are modular and form essentially identical structures in various folded RNAs (27–30), will likely often exhibit energetic independence and allow ρ analysis to be carrier out for additional and more complex RNAs and RNA-protein complexes.

### The P14 and MC/MCR tertiary contacts form after the TL/TLR in ^alt^P5abc•E^ΔP5abc^ **assembly**

Having established that TL/TLR formation is rate-limiting for ^alt^P5abc•E^ΔP5abc^ assembly using U167C and G174A P5abc, we tested whether the other two contacts, P14 and the MC/MCR, form before or after the rate-limiting step. We tested these alternatives by determining ϕ values for their formation, with the simplest expectation of values of 1 or 0, which would correspond to formation of the contact prior to or subsequent to rate-limiting TL/TLR formation, respectively (see Figure 1B,C).

To obtain these ϕ values, we measured binding and dissociation of the same P5abc variants described above to E^ΔP5abc^ variants with point mutations to weaken P14 or MC/MCR (see Fig. 2). Point mutations in E^ΔP5abc^ that disrupt base pairs in P14 (see Fig. 2A, green) increased the *k*_off_ values by as much as 100-fold but had only small effects on *k*_on_ values, at most three-fold (Fig. 4A; Fig. S1; Fig. S2). Similarly, point mutations in the MCR increased the dissociation rate of U167C P5abc with little effect on *k*_on_ (Fig. 4B). Thus, the ϕ values for formation of both contacts are near zero (Fig. 4C; Fig. S2; see Methods), providing strong evidence for their formation after the rate limiting TL/TLR formation, as in Fig. 1C. Stated another way, weakening these interactions does not slow assembly appreciably, indicating that the interactions are not already formed in the rate-limiting step for assembly. As the mutations affect the overall equilibrium and the disassembly rate, they are formed in the fully assembled complex, but after the rate-limiting step for assembly; due to microscopic reversibility they must break prior to the rate-limiting step for disassembly.

**Figure 4.**
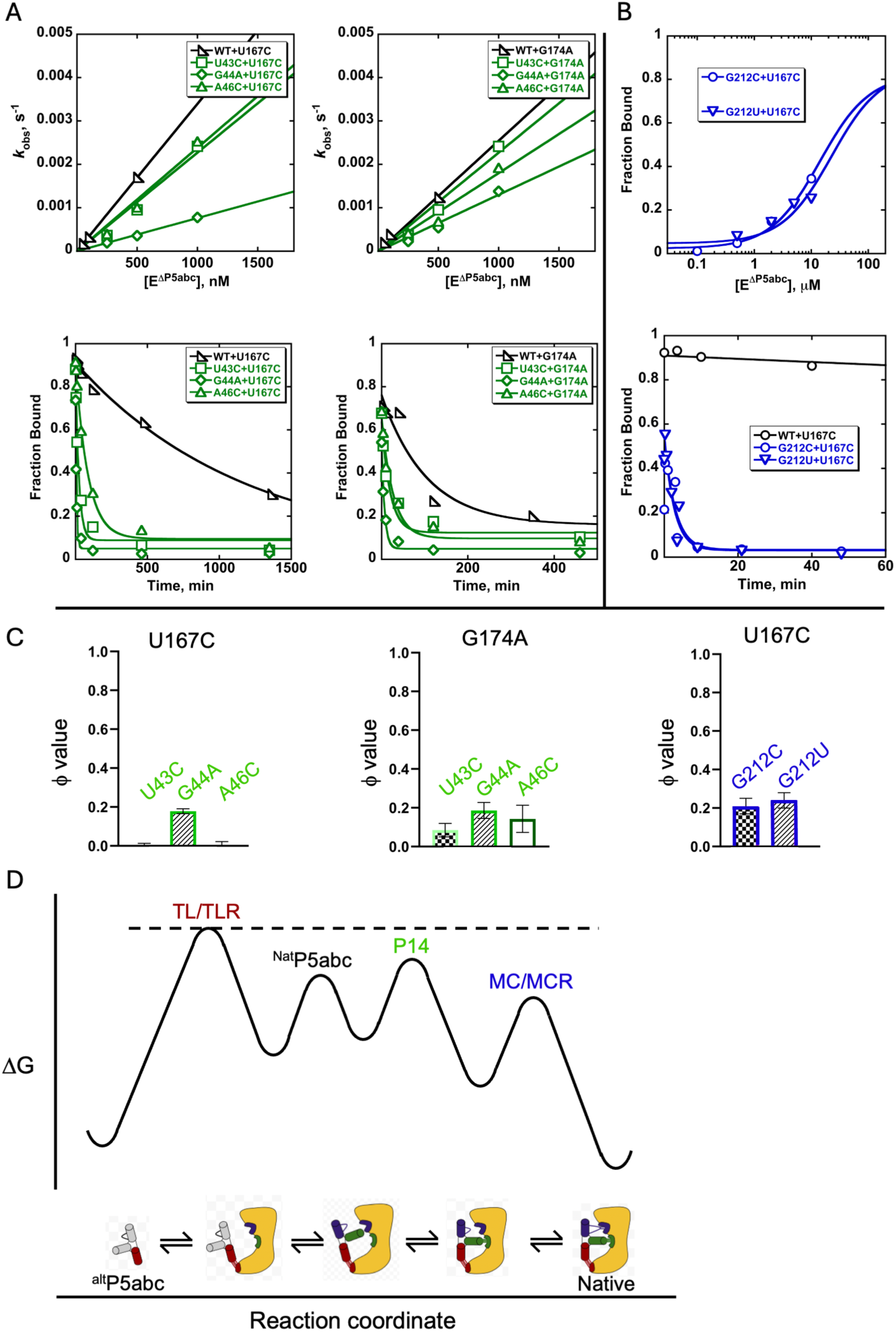
ϕ analysis of P14 and MC/MCR formation in the assembly pathways for P5abc variants with stabilized ^alt^P5abc. The mutants used are shown in Fig. 2A, and experiments were performed as shown in Fig. 3A,B. **A.** Top, concentration dependences of observed assembly rate constants for U167C P5abc RNA to WT E^ΔP5abc^ and the P14 mutants U43C, G44A, and A46C. Bottom, time courses for dissociation of U167C P5abc RNA from WT E^ΔP5abc^ and the P14 mutants U43C, G44A, and A46C. **B.** Top, concentration dependences of equilibrium binding of U167C P5abc RNA to the MC/MCR mutants G212C and G212U. Bottom, dissociation kinetics of U167C P5abc to the MC/MCR mutants. The effects of these mutations on equilibrium binding and dissociation rate constants were the same within error, indicating that the mutations do not decrease the binding rate constant (*k*_on_). **C.** Left & center panel: ϕ values for the P14 mutations U43C, G44A, and A46C in the context of P5abc variants that populate the ^alt^P5abc structure (U167C & G174A). Right panel: ϕ values for the MC/MCR mutants G212C and G212U in the context of the U167C P5abc RNA. **D.** Free energy profile for assembly of WT E^ΔP5abc^ with ^alt^P5abc. The TL/TLR contact forms first and is rate-limiting for the assembly process, giving ϕ values that match the corresponding ρ values (Fig. 3). We infer that the next step is the rearrangement of P5abc to ^Nat^P5abc, which results in native structure in the P5abc loops that form P14 and the MC/MCR. Because P14 and MC/MCR form subsequently, after the rate-limiting step, we cannot distinguish which one forms earlier or whether pathways with either are followed to similar extents. For simplicity, P14 is shown forming first.

Together these data indicate that the TL/TLR forms first and is rate-limiting for the overall assembly process (*i.e.*, the highest free energy barrier), and then P14 and the MC/MCR form, although we cannot distinguish their order of formation (Fig. 4D). For assembly to proceed downhill from this highest barrier, at least one subsequent contact must form before the TL/TLR dissociates (*k*_off,predicted_ = 1 s^−1^ from measurements of the isolated contact (6)); otherwise the TL/TLR would repeatedly form and dissociate without productive folding. The P5abc components of both the P14 and MC/MCR contacts are not properly formed in ^alt^P5abc but are present in the final complex (21, 26), indicating that ^alt^P5abc must rearrange to ^Nat^P5abc for either tertiary contact to form. Thus, this conformational transition must itself occur with a rate constant of >1 s^−1^. The secondary structure in P5c exchanges on the timescale of milliseconds (25), and our data indicate that the native secondary structure must be rapidly trapped by tertiary structure formation, allowing the entire transition to proceed within the ∼1 s lifetime of the intermediate.

### ρ analysis reveals a different folding pathway for WT P5abc RNA

We tested whether wild-type P5abc assembly with E^ΔP5abc^ follows a pathway with rate-limiting TL/TLR formation occurring first, as for ^alt^P5abc, or whether the differences in P5abc result in changes in the order of tertiary contact formation and/or the rate-limiting step. Unlike the P5abc mutants described above, wild-type P5abc exists predominantly in the native structure (^Nat^P5abc) even in the absence of E^ΔP5abc^ (see Fig. 2D). As a further test of our models, we also assayed the P5abc variant G176A, which, like wild-type P5abc, predominantly forms ^Nat^P5abc in the absence of E^ΔP5abc^ (20).

The E^ΔP5abc^ TLR point mutations, A225U and A226U, produced equilibrium effects for assembly with WT P5abc that were similar to the effects of the same mutations on the TL/TLR in isolation (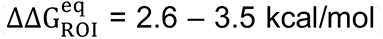), just as they did for assembly with the ^alt^P5abc-favoring mutants (U167C and G174A) described above (Fig. 5A; Fig. S3; Fig. S4). This finding further highlights the energetic independence common to RNA strctures and the potential broad applicability of ρ analysis and the reconstitution model (6, 19).

**Figure 5.**
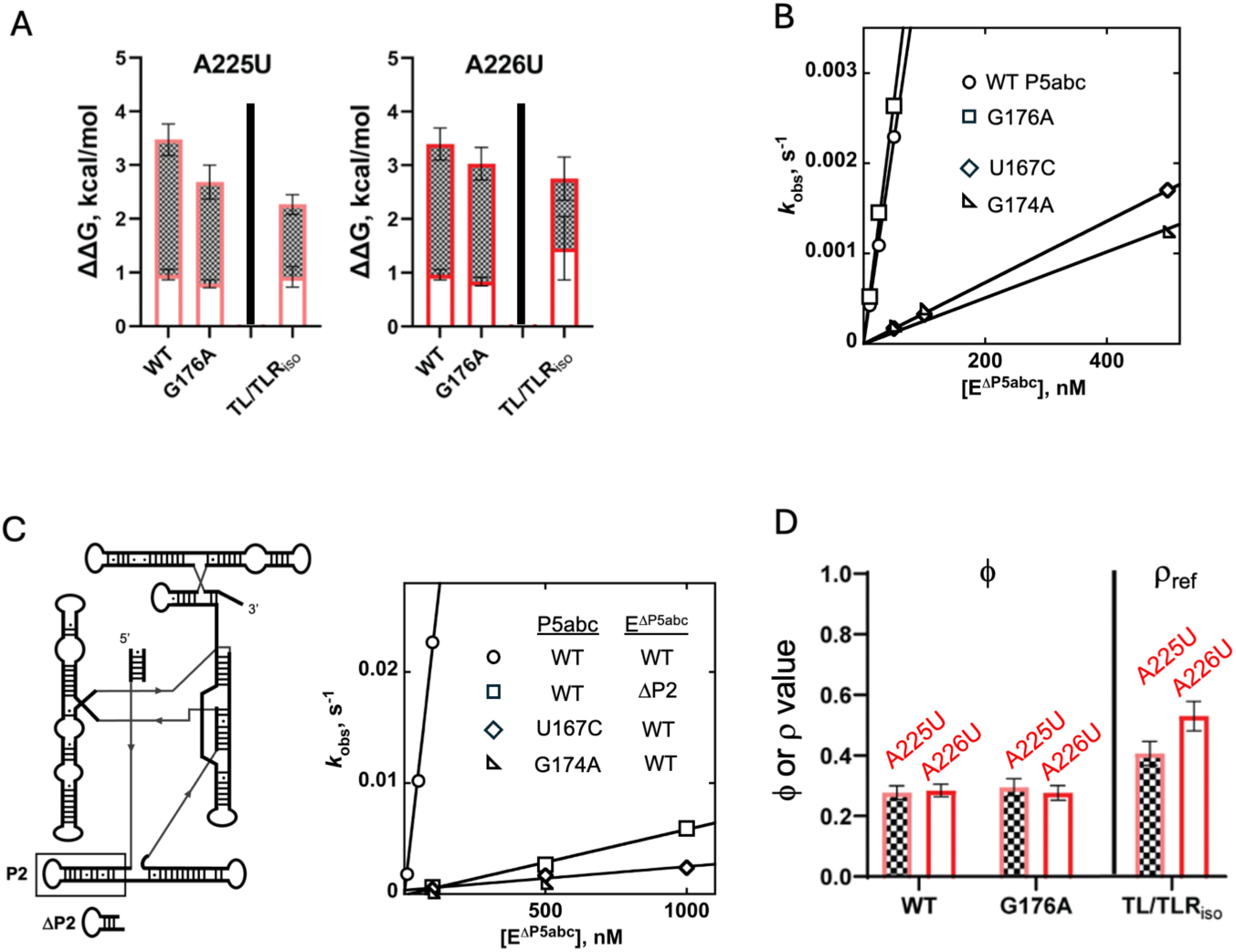
Analysis of TL/TLR and P14 formation in E^ΔP5abc^ assembly with WT P5abc and G176A P5abc, a variant that disrupts a base pair in ^alt^P5abc and, like WT P5abc, populates the native conformation (^Nat^P5abc). Experiments were performed as shown in Fig. 3A,B. **A.** Energetic effects of the TL/TLR mutations (A225U and A226U) on assembly (open bars) and disassembly (hatched bars) of WT P5abc RNA and the indicated variants with E^ΔP5abc^. The effects are shown for the same mutations in the context of isolated formation of the TL/TLR contact (from (6)). **B.** Concentration dependences of observed binding rate constants for E^ΔP5abc^ with for P5abc variants favoring the native conformation (^Nat^P5abc; WT and G176A). The analogous data for mutants favoring the alternative P5abc conformation (^alt^P5abc; U167C and G174A) are reprinted from Fig. 3 for comparison. The binding rate constant for ^Nat^P5abc, 6 X 10^4^ M^−1^ s^−1^, is substantially larger than the expected rate constant of ∼10^3^ M^−1^ s^−1^ for TL/TLR formation based on measurements of dissociation rate constant (∼3 s^−1^; (6, 19, 32)) and equilibrium dissociation constant under similar conditions (*K*_D_ *=* 3 mM; (31)). **C.** Left, secondary structure of the E^ΔP5abc^ construct and a schematic of the truncated stem in the ΔP2 variant. Right, the concentration dependences of observed binding rate constants; eliminating the ability to form P14 (ΔP2) reduces the binding rate constant for WT P5abc to levels similar to those for variants that populate ^alt^P5abc. (To facilitate measurements of the weakened complex resulting from the loss of P14, measurements were made at a lower temperature, 25 °C, and in the absence of added KCl; reactions of the WT complex were performed under the same conditions for comparison.) **D.** ϕ values for mutations in the TL/TLR contact in E^ΔP5abc^ (A225U, A226U; hatched and solid bars respectively) for association with ^Nat^P5abc (WT and G176A). These values are smaller than the ρ values for these mutations in the isolated TL/TLR (ρ_ref_), shown at right.

Turning to kinetics, we found that wild-type P5abc assembles with E^ΔP5abc^ 30-fold faster than ^alt^P5abc (Fig. 5B, (20)) and faster than the estimated association rate constant for an isolated TL/TLR (6, 31)). Thus, TL/TLR formation does not occur first in the wild-type P5abc assembly pathway. If it did, assembly could be no faster than this step; instead, the faster assembly indicates that a different, faster step must occur first, with subsequent TL/TLR formation.

We next tested whether the rapid assembly of WT P5abc is a result of rapid P14 formation, preceding TL/TLR formation. The simplest prediction of this model is that removal of P14 would slow assembly. To carry out this test we generated a E^ΔP5abc^ construct in which the P2 helix is shortened to three base pairs and the loop sequence is changed, eliminating the ability to form P14 (Fig. 5C).^§^ The loss of an ability to form P14 resulted in a 100-fold decrease in the binding rate constant for WT P5abc (Fig. 5C; Fig. S5; Table S4). Thus, the rapid assembly of native ^Nat^P5abc•E^ΔP5abc^ strongly depends on P14. As the assembly rate constant in the absence of P14 is the same within error as those for the U167C and G174A ^alt^P5abc mutants, TL/TLR formation appears to become rate limiting when P14 cannot form or when a need for rearrangement slows its formation (6, 7, 31, 32).

We now turn to TL/TLR formation. While the overall thermodynamic effects of the A225U and A226U E^ΔP5abc^ TLR mutations were largely the same for wild-type P5abc as they were for the P5abc mutants (U167C and G174A) and the isolated TL/TLR, their *kinetic* effects were consistently different (Fig. 5A). Specifically, the decreases in the binding rate constant were smaller (by 2–5-fold) and the increases in dissociation rate constant were larger (by 2–10-fold) with wild-type P5abc than with TL/TLR_iso_. As a result, the resulting ϕ values of 0.24–0.30, shown in Fig. 5D, are significantly lower than the values for TL/TLR formation in isolation (ρ_ref_ = 0.4–0.6) and those for ^alt^P5abc•E^ΔP5abc^ assembly (ϕ = 0.5–0.6). These effects indicate that there are additional differences in the assembly pathway with wild-type P5abc.

Overall, the above results for ^Nat^P5abc•E^ΔP5abc^ assembly strongly suggest that P14 forms prior to the TL/TLR. When P14 cannot form (from either stabilization of ^alt^P5abc or mutation of L2), assembly slows and TL/TLR formation becomes rate limiting. Nevertheless, there is additional complexity for ^Nat^P5abc•E^ΔP5abc^ assembly. The ϕ values for TL/TLR formation during ^Nat^P5abc•E^ΔP5abc^ assembly are significantly greater than zero, indicating that the TL/TLR does not simply form in a step subsequent to a rate-limiting step of P14 formation in a linear assembly pathway. One possible model invokes the presence of competing pathways involving initial TL/TLR or P14 formation, but TL/TLR formation is too slow to compete significantly with the ∼100 fold faster P14 loop-loop formation. It is possible that P14 formation is rate-limiting for assembly with WT E^ΔP5abc^ but that the TLR mutations (A225U and A226U) sufficiently destabilize the TL/TLR so that its formation becomes rate limiting for the mutants. Such a threshold effect would result in ϕ being smaller than ρ_ref_ as is observed (Fig. S6). Alternatively, geometric constraints from P14 may alter the energetics of microscopic steps and nature of the transition state for subsequent TL/TLR formation, for example by increasing the lifetimes of partially bound states and altering the TL/TLR association landscape.

Together, the results lead to the model depicted in a cartoon landscape in Fig. 6. WT P5abc, which folds primarily to ^Nat^P5abc at equilibrium, assembles with the ribozyme core by preferential formation of P14 (thick arrow at right), followed by formation of the TL/TLR and MC/MCR contacts. A second pathway, with the TL/TLR formed first, is presumably available to ^Nat^P5abc, but this pathway receives minimal flux (estimated at 1%) because of the intrinsically slower formation of the TL/TLR contact. On the other hand, P5abc variants that populate ^alt^P5abc and face a large energetic penalty for transient formation of ^Nat^P5abc assemble by forming the TL/TLR contact first, in the rate-limiting step as revealed by ρ analysis, and then rearranging to ^Nat^P5abc and forming P14 and the MC/MCR contacts.

**Figure 6.**
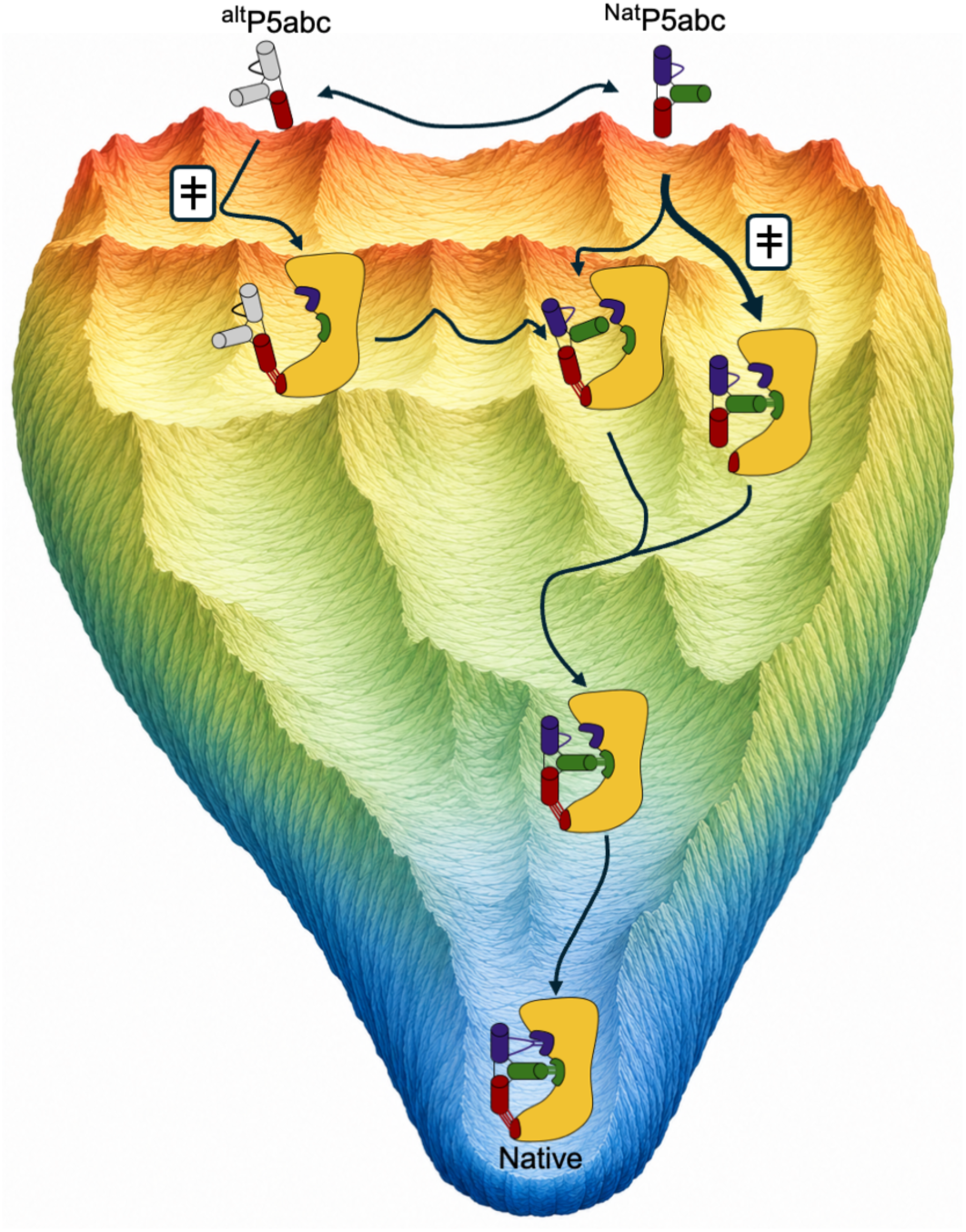
Model for P5abc folding and assembly landscape. The two folded conformations of P5abc in solution, ^Nat^P5abc and ^alt^P5abc, traverse the assembly landscape via distinct pathways. ^alt^P5abc assembles with initial formation of the TL/TLR contact, whereas ^Nat^P5abc assembles predominantly with initial formation of P14 (thick arrow at right). The pathways converge as ^alt^P5abc refolds to ^Nat^P5abc faster than it dissociates, and it then forms P14, while the pathway from ^Nat^P5abc continues with TL/TLR formation. In the simplest model and as suggested by the ρ analysis, the MC/MCR contact forms last along both of these pathways.

### Application of ρ analysis to DNA target binding by CRISPR-Cas12a

To further test and extend ρ analysis, we applied it to DNA target recognition by the CRISPR endonuclease Cas12a. Cas12a uses complementarity with a self-processed crRNA to bind a DNA target that also includes a protospacer adjacent motif (PAM), forming an R-loop between the crRNA and the target strand of the DNA and ultimately cleaving both DNA strands (Fig. 7A). Previous work has shown that Cas12a displays specificity against sequence differences (mismatches) between the crRNA and the DNA target (33), with mismatches typically giving penalties of 10-100-fold, but until now there was not a quantitative framework from which to assess these values or to consider, in general, the specificity levels that might be expected in this class of enzymes.

**Figure 7.**
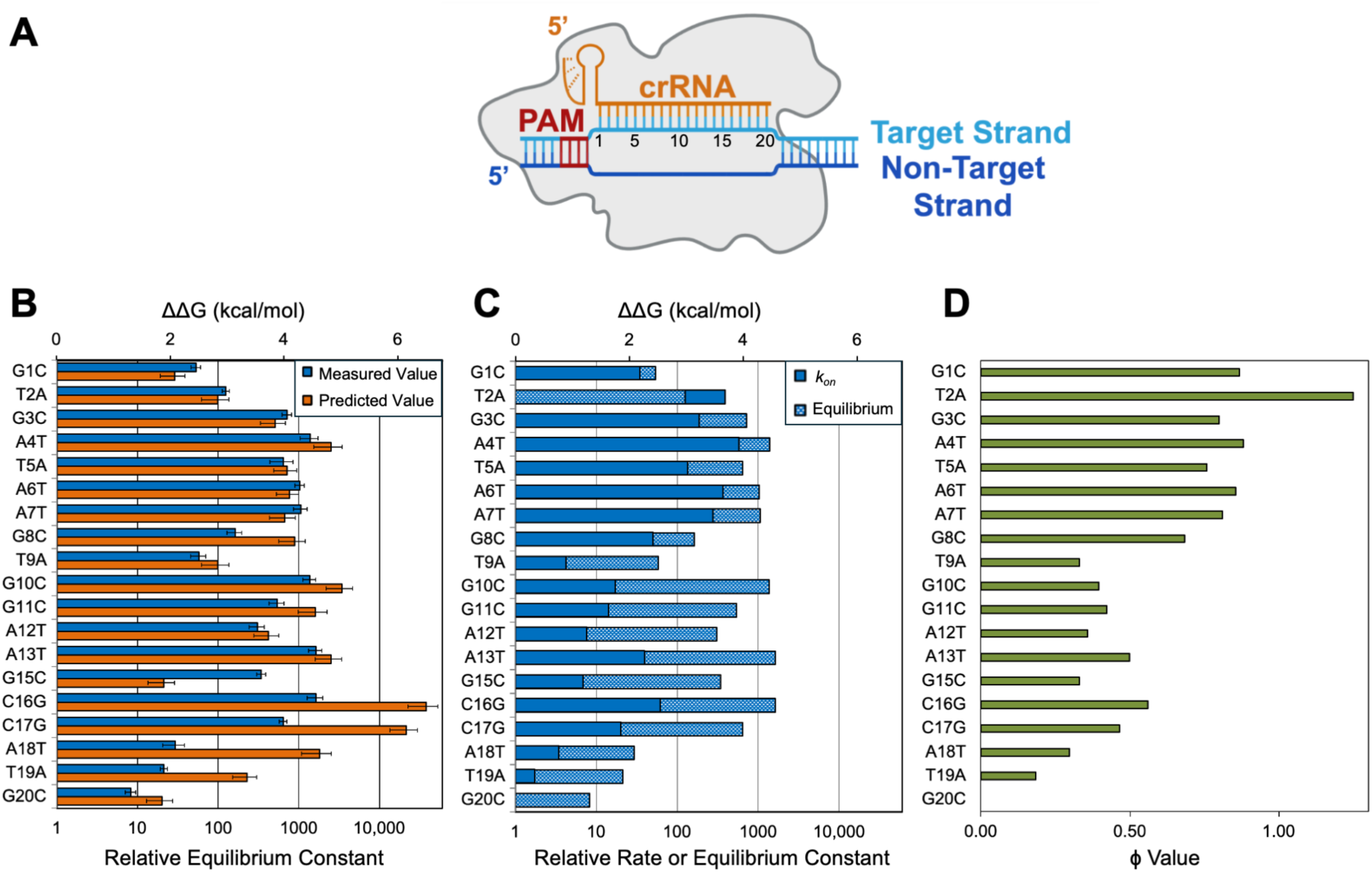
Effects of target mismatches on AsCas12a binding thermodynamics and kinetics. **A.** Schematic of Cas12a bound to a DNA target. The crRNA (orange) forms base pairs with the target strand of the DNA (cyan), displacing the non-target strand. The PAM sequence of the DNA is shown in red. **B.** Comparison of energetic penalties for mismatches between the DNA and crRNA, relative to binding of a DNA with full complementarity of the target strand to the crRNA at positions 1-20. Measurements are blue and nearest-neighbor predictions are orange (see Methods and Fig. S8 for details on how predicted values were calculated). Note that the sequences of both strands of the DNA were changed at a single position, such that complementarity was maintained between the DNA target and non-target strands, but complementarity with the crRNA was lost. Nearest-neighbor predictions at each position consider both the sequence change in the R-loop and that in the dsDNA. Labels at left indicate the sequence change in the non-target strand. The x-axis (log scale) gives the destabilization as a relative equilibrium binding constant. **C.** Comparison of energetics and kinetics from mismatches. For each position, the effect on binding kinetics is shown in solid blue and the overall equilibrium effect is shown in hatched blue. The corresponding ϕ values are shown at the right. **D.** ϕ values for mismatches at each position.

We therefore applied the principles of ρ analysis to an extensive existing dataset of AsCas12a binding to DNA target sequences *in vitro* (33). We first compared the equilibrium free energy penalties for each mismatch at each position within the R-loop (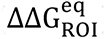), obtained from measuring binding and dissociation rate constants, to the penalties calculated from nearest-neighbor predictions for RNA:DNA duplexes (Fig. 7B). Overall, there was substantial agreement between the measured and predicted values, highlighting the modular energetics of base pairs and indicating a general value of nearest-neighbor rules as a framework for calculating 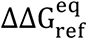 values for mutations that disrupt base pairing. It is especially striking that these rules largely hold even in the context of the Cas12a protein, which associates with both strands and forms a channel in which the R-loop lies.

For Cas12a, while most of the interactions followed expectations from nearest neighbor energetics, deviations were also observed. In particular, mismatches at the PAM-distal end of the R-loop resulted in smaller penalties than predicted. Given the prior evidence for a conformational change associated with formation of PAM-distal base pairs that activates nuclease activity (34, 35), the simplest model is that the energetic difference represents the cost, or “payment” for this conformational change. Intriguingly though, the ΔΔG values at these positions are not constant, suggesting that different mismatched states may be favored on Cas12a than in free helices. There are also small deviations from the nearest neighbor predictions at some internal sites. These differences may represent conformational distortions of the protein-bound base pairs that prevent them from attaining their thermodynamically most stable states and may be associated with additional conformational changes during R-loop propagation within the protein site.

Because dissociation of Cas12a is extremely slow, relative to subsequent DNA cleavage, specificity is determined during binding and is thus dependent on the kinetics of R-loop formation. We used the energetic penalties to calculate ϕ values as 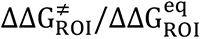 (Fig. 7C,D; Fig. S7; Eq. 1). The ϕ values are largest at the PAM-proximal end of the R-loop and decrease progressively toward the distal end, mirroring the direction of base pair formation in the propagating R-loop (34–39): base pairs formed early are present in the transition state to a greater extent than those formed later. Nevertheless, the ϕ values remain substantially above zero across most of the R-loop. This observation suggests that the transition state for R-loop formation is late and that partially formed R-loops can reversibly sample downstream base-pairing interactions prior to commitment.

In this framework, the thermodynamic mismatch penalties define an upper limit on the specificity that is achievable, as they represent the full energetic difference between binding to the matched vs the mismatched sequence. In that contect, the observed ϕ values report on how much of that potential is realized in kinetic discrimination by Cas12a. The persistence of nonzero ϕ values throughout the R-loop indicates a late transition state and provides a mechanistic basis for Cas12a’s substantial discrimination against mismatches across the extended target region, despite the fact that binding is rate-limiting. Importantly, this late transition state differs from that for formation of a simple duplex in solution, which occurs with only 2–3 base pairs formed (40). We expect that this upper limit on specificity will be general for CRISPR-Cas enzymes that target RNA as well as those that target DNA (41), and it may hold more broadly for proteins that use complementarity with a guide RNA to target a DNA or RNA with sequence specificity.

## CONCLUSIONS & IMPLICATIONS

We have developed ρ analysis for quantitative analysis of RNA conformational processes including folding, complex assembly, and function. In essence, ρ analysis compares experimentally-determined ϕ values for a complex assembly process to the corresponding values for the same mutations for formation of the isolated contact. As a result, ρ analysis incorporates a uniquely powerful reference set that (1) assesses the modularity of RNA folding processes and (2) diagnoses the order of conformational steps, the rate-limiting step or steps, and whether multiple pathways are followed. Our results with the *Tetrahymena* group I intron ribozyme provide further evidence for modular energetics in RNA tertiary assembly and demonstrate changes in RNA assembly pathways from mutations. Our results with CRISPR-Cas12a demonstrate that ρ analysis can be used to dissect changes in RNA secondary structure during function, revealing the mechanisms and limits of DNA target specificity. Together the results establish ρ analysis as a powerful quantitative tool to dissect RNA folding and assembly pathways.

Here we demonstrated that ρ analysis is applicable for the multi-step assembly of the *Tetrahymena* group I intron from RNA components. ρ analysis is also applicable to functions that involve conformational rearrangements, as is common in RNA-mediated processes including pre-mRNA splicing by the spliceosome, translation by the ribosome, telomere maintenance by telomerase, and DNA and RNA targeting by Cas9 and related CRISPR endonucleases. Importantly, events involving different RNA targets may have different rate-limiting steps and, for events such as alternative splicing, different product-determining steps. Thus, studies with common modifications may help distinguish different control steps for different RNAs.

Further, as shown by our analysis of Cas12a, ρ analysis can be used analogously to assess secondary structure formation and rearrangements, with implications for the process of DNA binding by Cas12a and expected limits on the specificity against mismatches with DNA sequences. It is likely that there will prove to be a broader range of applications involving RNA secondary structure than tertiary structure, because more functional RNAs form secondary structure and because there are extensive thermodynamic rules for base pairs and other secondary structure elements. Examples where forming and changing secondary structure include control via riboswitches and steps in viral RNA folding and packaging. Further, as ρ analysis directly monitors the effects of mutations in secondary and tertiary structure elements, these effects provide a “signature”. By measuring how this signature is altered by RNA modifications and specific RNA binding proteins, one can determine whether —and how— these features alter RNA folding and assembly processes.

For the formation of complex RNAs and RNA-proteins assemblies it is typically not possible to measure effects on overall stability, precluding traditional ϕ analysis. In contrast, the comparisons to the energetics of external isolated RNA elements in ρ analysis will allow these complex and important RNA-mediated processes to be studied in depth. Further, for changes in secondary structure, our results show that Nearest Neighbor predictions provide a good approximation of energetic penalties (see Fig. S7), allowing ρ analysis to be used even in the absence of measurements on isolated RNA elements. Finally, ρ analysis is also readily carried out *in vivo*. In particular, ρ analysis may aid in dissecting complex cellular decisions such as alternative pre-mRNA splicing to determine the features and interactions responsible for favoring one splicing outcome over another.

## MATERIALS AND METHODS

### RNA Preparation

RNAs were generated by *in vitro* transcription using T7 RNA polymerase and DNA plasmid templates, which were digested with BsaI for P5abc RNAs or ScaI for E^ΔP5abc^ RNAs, under conditions as previously described (42). P5abc RNAs were ^32^P-labeled by incubating RNA with Shrimp Alkaline Phosphatase (New England Biolabs) and then with T4 polynucleotide kinase (New England Biolabs). Radiolabeled RNAs were purified by non-denaturing 12% PAGE. Unlabeled ribozyme core variants and P5abc were purified using an affinity column (Qiagen).

### Assembly and disassembly kinetics measurements

Rate constants were measured using a pulse-chase protocol. RNAs were pre-folded as previously described (20). Unless otherwise indicated, reactions conditions were 50 mM Na-MOPS pH 7.0, 10 mM MgCl_2_, and 100 mM KCl at 37 °C. For assembly rate measurements, trace amounts of ^32^P-labeled P5abc (wild-type or variants) were mixed with varying concentrations of wild-type or variant E^ΔP5abc^, and reaction aliquots were quenched at various times by increasing the Mg^2+^ concentration to 50 mM, adding a 10-fold excess of unlabeled P5abc relative to E^ΔP5abc^, and placing the quenched aliquot on ice. The high Mg^2+^ concentration and low temperature ensured that the assembled complex did not dissociate prior to resolution by gel electrophoresis, and the excess unlabeled P5abc blocked further complex assembly. Aliquots reflecting time zero were generated by premixing the labeled and unlabeled P5abc and then adding the mixture to E^ΔP5abc^. Complexes were resolved by 12% native PAGE and quantified using phosphorimager analysis.

For disassembly rate measurements, the complex was formed between radiolabeled P5abc and E^ΔP5abc^ (typically 3–5 µM E^ΔP5abc^) and then diluted 10-fold with an excess of 10-fold unlabeled P5abc. Reaction aliquots were quenched by adding buffer on ice and increasing the Mg^2+^ concentration to 50 mM and were processed by PAGE as above. The fraction of labeled P5abc that was bound at the start of each time course (a zero time point) was measured by mixing the bound complex with unlabeled chase and quench simultaneously. The expected endpoints of dissociation curves were determined by premixing the labeled and unlabeled P5abc and adding the mixture to E^ΔP5abc^. For variant combinations with very low binding affinity (the MC/MCR mutants), assembly rate constants could not be determined directly due to precipitation of E^ΔP5abc^ at high concentrations. Instead, an apparent equilibrium constant was measured by incubating the complex at varying concentrations of E^ΔP5abc^, and the assembly rate constant was calculated from the relationship between the equilibrium and dissociation rate constants.

An incubation time appropriate to reach equilibrium was determined from the measured dissociation rates of these unstable variants (incubations ranged from 3-16 hr).

### Data Processing

The fraction of bound radiolabeled P5abc over time was determined by quantifying the intensity of the labeled complex relative to the total intensity at each time point. Progress curves were fit to single exponential functions to determine the association and dissociation rate constants. Uncertainties reported for rate constants reflect the SEM from at least two replicate measurements. ϕ values for each combination of P5abc and E^ΔP5abc^ variants were determined by assessing the relative energetic effect of E^ΔP5abc^ mutations on the binding equilibrium and dissociation kinetics: ϕ = (ΔΔ*G*^‡^/ΔΔ*G*^Eq^) = log(*k*_on,mut_ / *k*_on,WT_) / log((*k*_on,mut_ × *k*_off,WT_) / (*k*_on,WT_ × *k*_off,mut_)).

### CRISPR-Cas12a analysis

Kinetics measurements of AsCas12a binding and dissociation from DNA target D were taken from ref. (33). Equilibrium constants for Cas12a binding to DNA targets with and without mismatches with the crRNA were calculated from the corresponding binding and dissociation rate constants (*K*_D_ = *k*_off_ / *k*_on_). Predicted values were calculated using parameters defined from experiments for DNA duplexes (43) and RNA-DNA hybrid duplexes (44, 45) under standard conditions at 37 °C. For each mismatch, the predicted free energy penalty was defined as the difference between the predicted free energy change for formation of the matched vs mismatched R-loop from the corresponding DNA duplex (Fig. S8).

## ACKNOWLEDGMENTS

We thank Chet Graswich and John Shin for figure preparation. This research was supported by NIH grants R35 GM131777 to R.R. and R01 GM132899 to D.H.

## Abbreviations

E^ΔP5abc^,: P5abc-deleted *Tetrahymena* ribozyme
EMSA,: electrophoretic mobility shift assay
MC/MCR,: metal core/metal core receptor tertiary contact
PAGE,: polyacrylamide gel electrophoresis
TL/TLR,: tetraloop/tetraloop receptor tertiary contact

^^ 

^^ 

## Footnotes

‡ The Reconstitution Model is readily extended to incorporate secondary structure transitions prior to tertiary structure formation and the stable formation of certain secondary structure elements only after particular tertiary contacts are made.

§ We also blocked P14 formation by mutating the L2 or L5c loops to UUCG while maintaining the stem lengths. These mutations each reduced the *k*on value to ∼10 M^−1^ s^−1^, 100-fold slower than expected for assembly through initial, rate-limiting TL/TLR formation (data not shown). This result suggests that the simultaneous presence of full-length P2 and L5c inhibits assembly if the loops cannot form base pairs, perhaps via steric clashes.

