## Supplemental tables and figures for "Rho (ρ) Analysis to Dissect RNA Folding and Assembly Pathways"

**Supplemental Information**

| **P5abc Variant** | $E^{\Delta P5abc}$ | U43C | G44A | A46C | A225U | A226U | G212C | G212U |
|---|---|---|---|---|---|---|---|---|
| wild-type | 6.3(±0.5)×10$^4$<br>(1) | 1.0(±0.1)×10$^4$<br>0.16 (±0.02) | 9.3(±0.2)×10$^3$<br>0.15 (±0.01) | 1.8±(0.2)×10$^4$<br>0.29 (±0.03) | 1.3(±0.1)×10$^4$<br>0.21 (±0.02) | 1.3(±0.1)×10$^4$<br>0.20 (±0.02) | 2.7×10$^4$*<br>0.42* | 6.7×10$^4$*<br>1.1* |
| G176A | 5.5(±0.3)×10$^4$<br>(1) | 2.2(±0.3)×10$^4$<br>0.38 (±0.05) | 9.7(±0.2)×10$^3$<br>0.17 (±0.01) | 2.2(±0.2)×10$^4$<br>0.39 (±0.03) | 1.5(±0.1)×10$^4$<br>0.27 (±0.02) | 1.4(±0.1)×10$^4$<br>0.24 (±0.02) | N.D. | N.D. |
| U167C | 2.2(±0.2)×10$^3$<br>(1) | 2.3(±0.2)×10$^3$<br>1.1 (±0.1) | 7.7(±0.2)×10$^2$<br>0.36 (±0.02) | 2.3(±0.2)×10$^3$<br>1.1 (±0.1) | 2.3(±0.3)×10$^2$<br>0.11 (±0.02) | 2.3(±0.3)×10$^2$<br>0.11 (±0.02) | 4.8×10$^2$*<br>0.22* | 3.2×10$^2$*<br>0.15* |
| G174A | 2.3(±0.2)×10$^3$<br>(1) | 1.8(±0.2)×10$^3$<br>0.73 (±0.07) | 1.3(±0.1)×10$^3$<br>0.55 (±0.06) | 1.8(±0.2)×10$^3$<br>0.76 (±0.09) | 1.3(±0.2)×10$^2$<br>0.053 (±0.01) | 2.5(±0.3)×10$^2$<br>0.11 (±0.02) | N.D. | N.D. |

**Table S1:** Association rate constants. In each cell, the top line indicates the second-order rate constant for the indicated combination of P5abc and ribozyme core (in units of $M^{-1}$ $s^{-1}$) and the second line indicates the rate relative to the corresponding value for the wild-type $E^{\Delta P5abc}$ core. Values that were calculated from equilibrium and dissociation rate constants are indicated with an (*). Uncertainties reflect the standard error from at least two independent measurements except for the double mutants, for which rate constants were measured only once. For these mutants, we used a conservative error estimate of 25%. N.D., not determined.

| **P5abc Variant** | $E^{\Delta P5abc}$ | U43C | G44A | A46C | A225U | A226U | G212C | G212U |
|---|---|---|---|---|---|---|---|---|
| wild-type | $6.5(\pm0.8)\times10^{-6}$<br>(1) | $5.8(\pm0.3)\times10^{-5}$<br>9.2 (±1.3) | $1.1(\pm0.3)\times10^{-3}$<br>170 (±50) | $3.2\pm(0.7)\times10^{-5}$<br>4.9 (±1.3) | $4.0(\pm0.1)\times10^{-4}$<br>62 (±18) | $3.5(\pm0.8)\times10^{-4}$<br>54 (±16) | $7.5(\pm0.2)\times10^{-3}$<br>1,200 (±160) | $6.0(\pm0.2)\times10^{-3}$<br>930 (±130) |
| G176A | $6.3(\pm1.8)\times10^{-6}$<br>(1) | $7.5(\pm0.8)\times10^{-5}$<br>12 (±4) | $4.3(\pm0.7)\times10^{-4}$<br>67 (±22) | $4.0(\pm0.5)\times10^{-5}$<br>6.2 (±2.0) | $1.4(\pm0.2)\times10^{-4}$<br>22 (±7) | $2.3(\pm0.3)\times10^{-4}$<br>36 (±11) | N.D. | N.D. |
| U167C | $2.5(\pm0.5)\times10^{-5}$<br>(1) | $8.5(\pm1.3)\times10^{-4}$<br>34 (±9) | $3.2(\pm0.8)\times10^{-3}$<br>130 (±45) | $2.5(\pm0.5)\times10^{-4}$<br>10 (±3) | $2.5(\pm0.2)\times10^{-4}$<br>10 (±2) | $3.2(\pm0.3)\times10^{-4}$<br>13 (±3) | $6.5(\pm1.3)\times10^{-3}$<br>260 (±76) | $8.0(\pm1.2)\times10^{-3}$<br>330 (±85) |
| G174A | $1.8(\pm0.8)\times10^{-4}$<br>(1) | $2.5(\pm1.2)\times10^{-3}$<br>13 (±8) | $2.2(\pm0.3)\times10^{-3}$<br>11 (±5) | $7.8(\pm1.3)\times10^{-4}$<br>4.2 (±1.9) | $1.0(\pm0.3)\times10^{-3}$<br>5.4 (±2.8) | $8.5(\pm1.5)\times10^{-4}$<br>4.5 (±2.1) | N.D. | N.D. |

**Table S2:** Dissociation rate constants. In each cell, the top line indicates the first order rate constant for the indicated combination of P5abc and ribozyme core (in units of $s^{-1}$) and the second line indicates the rate relative to the corresponding value for the wild-type $E^{\Delta P5abc}$ core. N.D., not determined.

| P5abc Variant | $E^{\Delta P5abc}$ | U43C | G44A | A46C | A225U | A226U | G212C | G212U |
|---|---|---|---|---|---|---|---|---|
| wild-type | $1.0(\pm0.2)\times10^{-10}$<br>(1) | $5.7(\pm0.6)\times10^{-9}$<br>56 (±10)<br>2.4 (±0.1) | $1.2(\pm0.3)\times10^{-7}$<br>1,200 (±340)<br>4.2 (±0.2) | $1.7(\pm0.4)\times10^{-9}$<br>17 (±5)<br>1.7 (±0.2) | $3.1(\pm0.8)\times10^{-8}$<br>300 (±92)<br>3.4 (±0.2) | $2.7(\pm0.7)\times10^{-8}$<br>270 (±80)<br>3.3 (±0.2) | $2.8(\pm0.1)\times10^{-7}$<br>2,700 (±420)<br>4.7 (±0.1) | $9.0(\pm0.4)\times10^{-8}$<br>880 (±140)<br>4.0 (±0.1) |
| G176A | $1.1(\pm0.3)\times10^{-10}$<br>(1) | $3.6(\pm0.6)\times10^{-9}$<br>32 (±11)<br>2.0 (±0.2) | $4.4(\pm0.7)\times10^{-8}$<br>390 (±130)<br>3.5 (±0.2) | $1.8(\pm0.3)\times10^{-9}$<br>16 (±5)<br>1.6 (±0.2) | $9.5(\pm1.2)\times10^{-9}$<br>84 (±27)<br>2.6 (±0.2) | $1.7(\pm0.2)\times10^{-8}$<br>150 (±50)<br>3.0 (±0.2) | N.D. | N.D. |
| U167C | $1.1(\pm0.3)\times10^{-8}$<br>(1) | $3.7(\pm0.7)\times10^{-7}$<br>32 (±9)<br>2.1 (±0.2) | $4.0(\pm1.2)\times10^{-6}$<br>350 (±130)<br>3.5 (±0.2) | $1.0(\pm0.2)\times10^{-7}$<br>9.2 (±3)<br>1.3 (±0.2) | $1.1(\pm0.2)\times10^{-6}$<br>92 (±25)<br>2.7 (±0.2) | $1.3(\pm0.2)\times10^{-6}$<br>120 (±33)<br>2.8 (±0.2) | $1.3(\pm0.3)\times10^{-5}$<br>1,200 (±350)<br>4.2 (±0.2) | $2.6(\pm0.4)\times10^{-5}$<br>2,300 (±600)<br>4.6 (±0.2) |
| G174A | $7.9(\pm3.4)\times10^{-8}$<br>(1) | $1.4(\pm0.6)\times10^{-6}$<br>18 (±11)<br>1.7 (±0.4) | $1.6(\pm0.3)\times10^{-6}$<br>21 (±10)<br>1.8 (±0.3) | $4.3(\pm0.8)\times10^{-7}$<br>5.5 (±2.6)<br>1.0 (±0.3) | $8.0(\pm2.8)\times10^{-6}$<br>100 (±57)<br>2.7 (±0.3) | $3.3(\pm0.7)\times10^{-6}$<br>42 (±20)<br>2.2 (±0.3) | N.D. | N.D. |

**Table S3:** Equilibrium constants. In each cell, the top line indicates the equilibrium dissociation constant ($K_d$) for the indicated combination of P5abc and ribozyme core (in units of M). The second line indicates the relative equilibrium constant to the corresponding value for the wild-type $E^{\Delta P5abc}$ core. The third line expresses the relative value as a ΔΔG value (in units of kcal $mol^{-1}$). N.D., not determined.

| P5abc Variant | WT $E^{\Delta P5abc}$ | $\Delta$P2 $E^{\Delta P5abc}$ |
|---|---|---|
| wild-type | $2.2(\pm 0.2)\times 10^5$<br>(1) | $2.6(\pm 0.5)\times 10^3$<br>(0.011) |
| U167C | $5.8(\pm 0.7)\times 10^3$<br>(0.026) | N.D. |
| G174A | $2.1(\pm 0.4)\times 10^3$<br>(0.010) | N.D. |

**Table S4:** Association rate constants with the $E^{\Delta P5abc}$ variant lacking P2 and therefore unable to form the P14 tertiary contact ($\Delta$P2). In each cell, the top line indicates the second-order rate constant for the indicated combination of P5abc and ribozyme core (in units of $M^{-1}$ $s^{-1}$) and the second line indicates the rate relative to the corresponding value for the wild-type $E^{\Delta P5abc}$ core and WT P5abc. The corresponding data are shown in Fig. 5C. Reaction conditions were 50 mM Na-MOPS, pH 7.0, and 10 mM $MgCl_2$ at 25 °C. N.D., not determined.

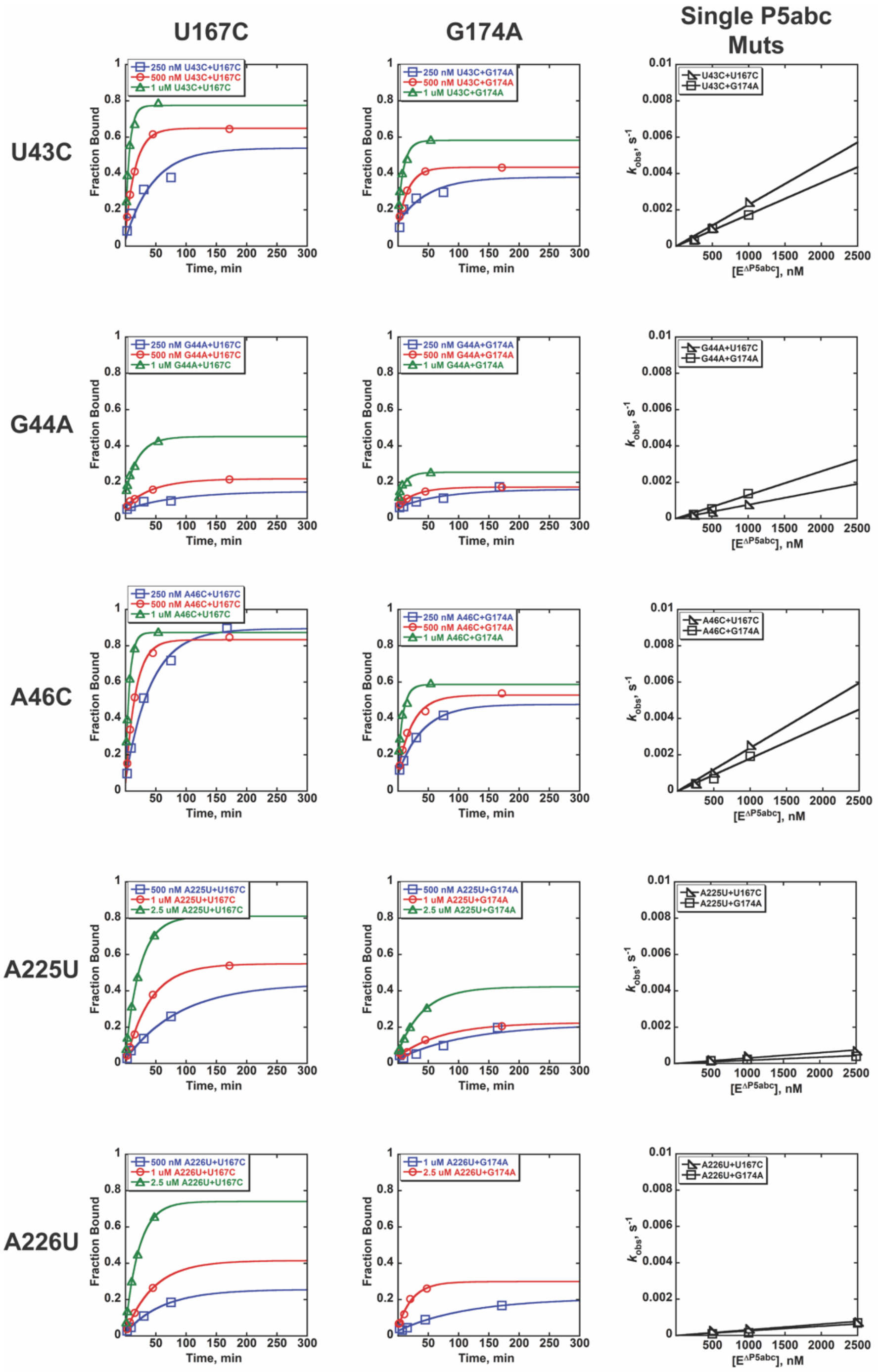


**Fig. S1.** Association kinetics of $E^{\Delta P5abc}$ mutants with P5abc variants that stabilize Alt P5abc, associating with $E^{\Delta P5abc}$ slowly and then rearranging to native structure (see Figure 2(b)). Measurements were quenched with a 5-fold excess of unlabeled wild-type P5abc relative to the concentration of $E^{\Delta P5abc}$. The fraction bound at t=0 is typically less than 20% because the unlabeled wild-type P5abc competes favorably against the labeled P5abc variants.

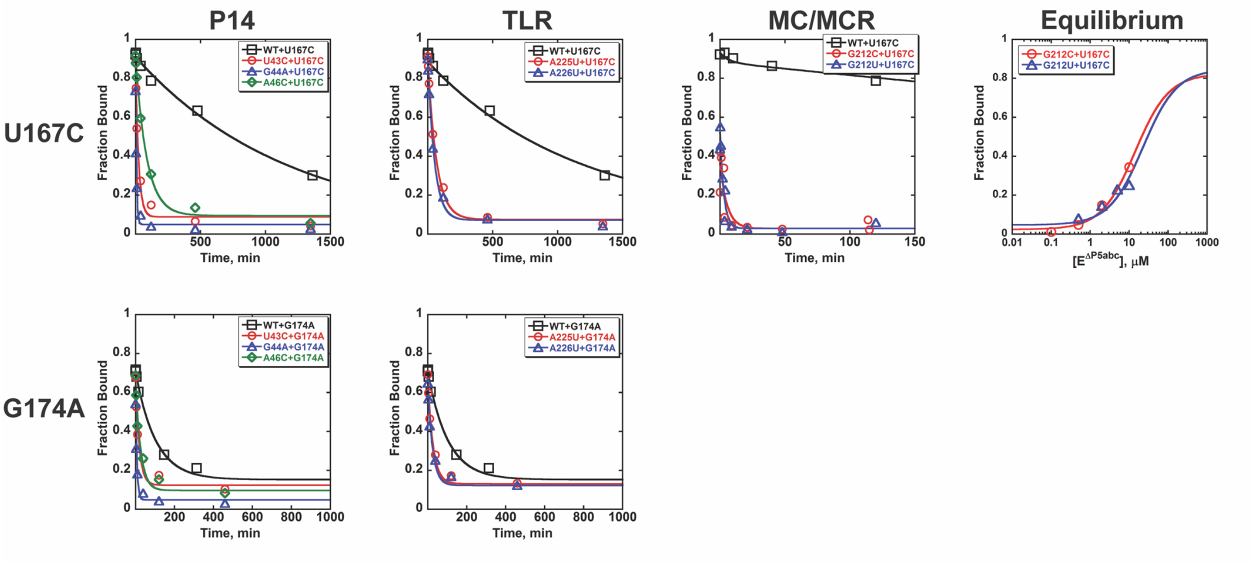


**Fig. S2.** Dissociation kinetics of tertiary contact mutants of $E^{\Delta P5abc}$ with P5abc mutants that stabilize the Alt P5abc structure. The right panel shows equilibrium binding curves for MC/MCR mutants. These curves are extrapolated to an endpoint that is observed for the wild-type $E^{\Delta P5abc}$ and respective mutant P5abc combination measured on the same day. The point mutations in $E^{\Delta P5abc}$ that weaken the MC/MCR contact (G212C and G212U) increased the dissociation rates by approximately 1000-fold (left panels) and weakened equilibrium binding by approximately the same amount (right panels). Equilibrium binding and dissociation measurements are used to calculate the $k_{on}$ values for these complexes, resulting in $\phi$ values <0.2 for WT P5abc (not shown) and a value of ~0.2 of U167C. Thus, MCR mutations that greatly destabilize the contact do not affect P5abc binding kinetics, with the large energetic penalty (4–5 kcal/mol) expressed largely or exclusively as an increased dissociation rate. These results suggest that the MC/MCR forms late in both the alternative and native pathways, after the rate-limiting transition state.

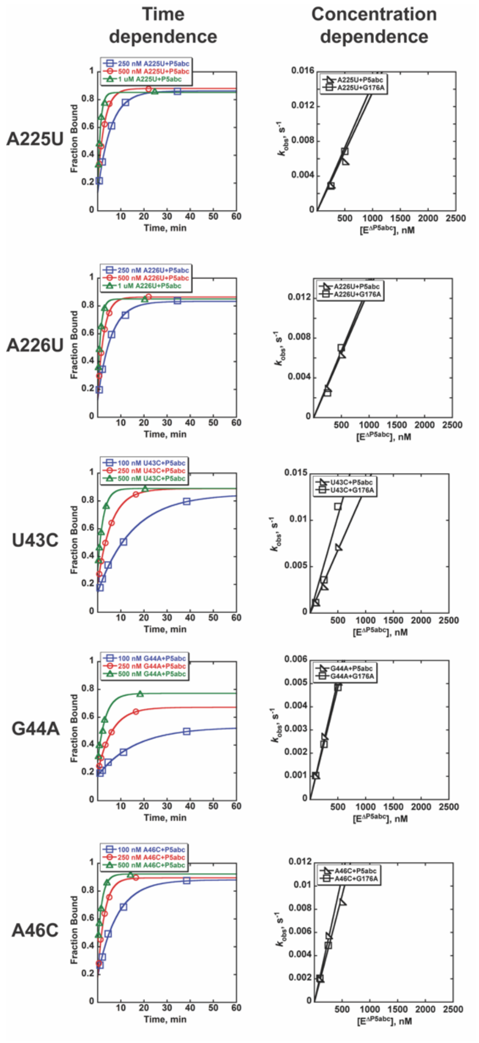


**Fig. S3.** Association kinetics of E$^{\Delta P5abc}$ mutants with WT P5abc and a P5abc variant that populates the native P5abc structure. Here and in subsequent supporting figures, measurements are shown from a given day, typically performed for a series of P5abc variants side by side. The rate constants from these measurements are within 2-fold of the values in Tables S1 and S2 (which reflect the averages of all measurements). The left panels show association of the indicated E$^{\Delta P5abc}$ mutant with the wild-type P5abc, and the right panels show dependences of the observed rate constant on the E$^{\Delta P5abc}$ concentrations for these reactions and for analogous reactions using the P5abc variant G176A, which also populates primarily the native P5abc structure. Reactions were quenched by adding 5-fold excess wild-type P5abc relative to E$^{\Delta P5abc}$ so that approximately 20% of the labeled P5abc is bound at t=0.

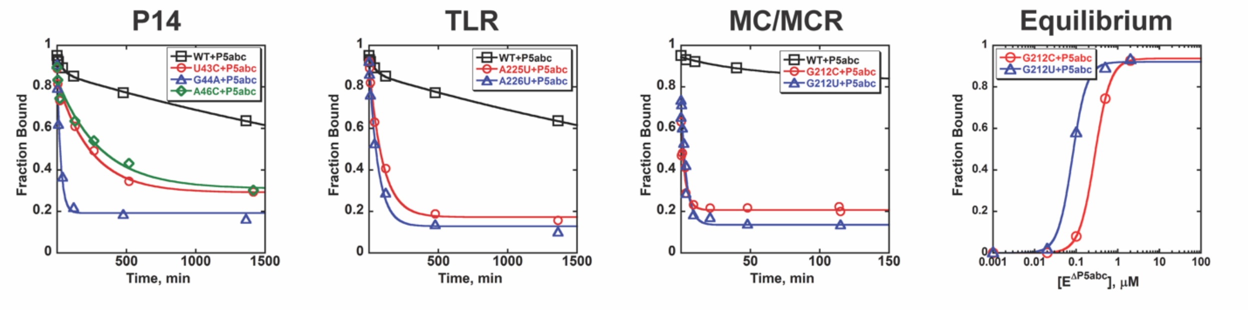


**Fig. S4.** Dissociation kinetics and equilibrium binding of $E^{\Delta P5abc}$ mutants from wild-type P5abc. For dissociation of wild-type $E^{\Delta P5abc}$, a rapid burst of dissociation is observed (~5%), which likely reflects binding by a fraction of misfolded or damaged RNA. The plot at the right shows equilibrium measurements of $E^{\Delta P5abc}$ binding by MC/MCR mutants of P5abc. These measurements were used with the measured $k_{off}$ values to calculate $k_{on}$ values for these mutants.

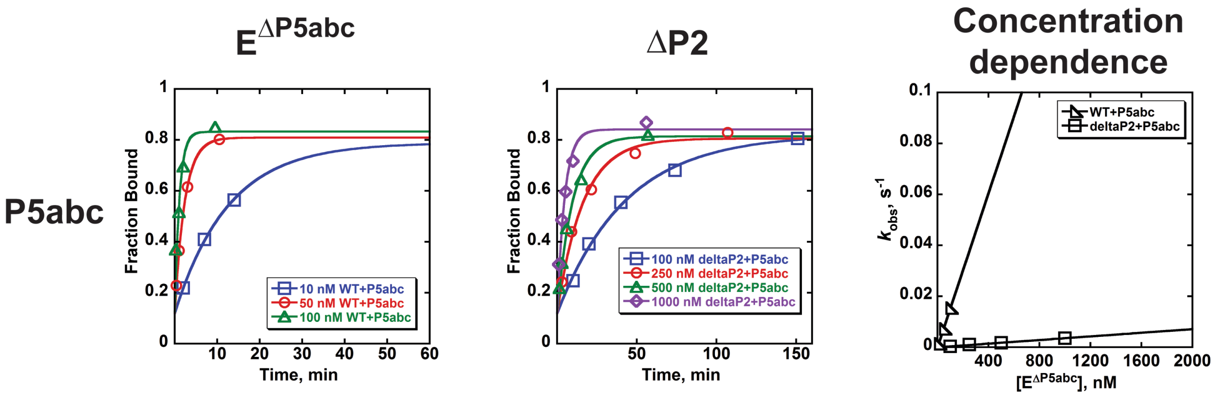


**Fig. S5.** Association kinetics of WT P5abc with the ΔP2 $E^{\Delta P5abc}$ mutant, with association of P5abc with the WT $E^{\Delta P5abc}$ shown under the same conditions for reference. These experiments were performed at 25 °C and in the absence of added KCl. Rate constants were $2.2 \times 10^5\ M^{-1}\ s^{-1}$ for association of P5abc with WT $E^{\Delta P5abc}$ and $5.8 \times 10^3\ M^{-1}\ s^{-1}$ for association with the ΔP2 $E^{\Delta P5abc}$ mutant.

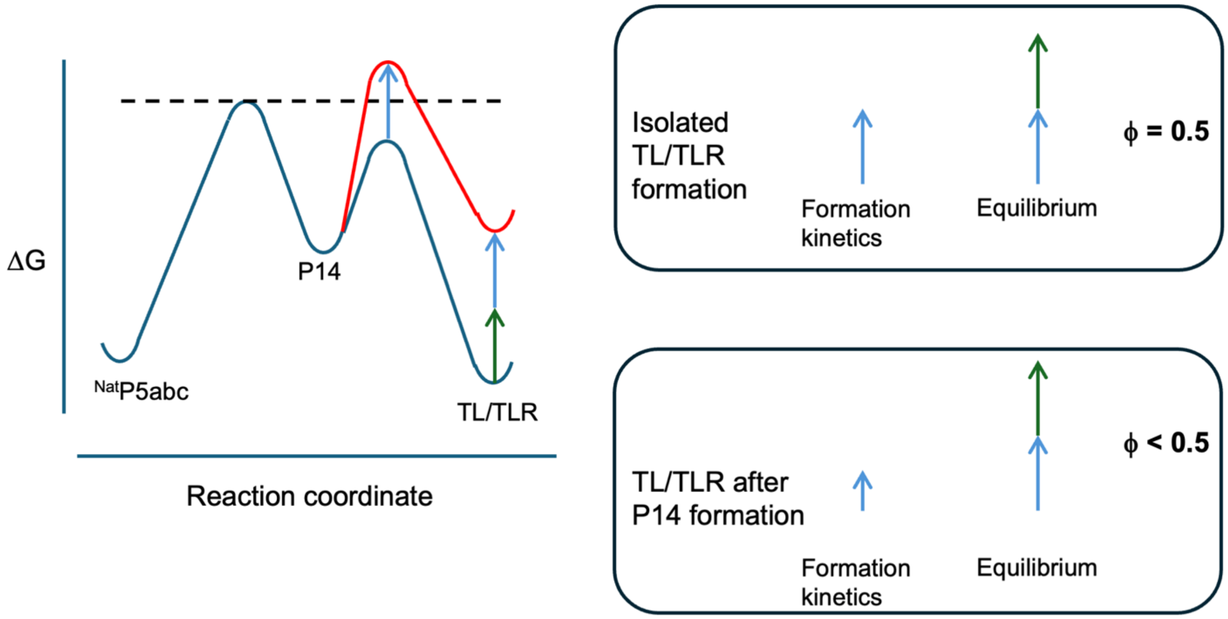


**Fig. S6.** Model for decreased $\phi$ value ($\phi < \rho_{ext}$) arising from a threshold effect. In this model, P14 forms first when $^{Nat}$P5abc binds to the WT $E^{\Delta P5abc}$ ribozyme (blue free energy profile). Upon mutation of the TLR, TL/TLR formation becomes rate-limiting for association (red curve). The blue and green arrows depict the free energy penalties for binding and dissociation, respectively, such that the equilibrium penalty is the sum of the two arrows as shown. For formation of the TL/TLR in isolation, the fraction of the free energy penalty that is expressed as a decrease in the binding rate constant is approximately 0.5 (top right box), but in the two-step process shown, the observed penalty on binding kinetics is smaller because only the portion that extends beyond the highest peak is observed (dashed line in free energy profile). Thus, the corresponding $\phi$ value is decreased, as shown in the box at the bottom right.

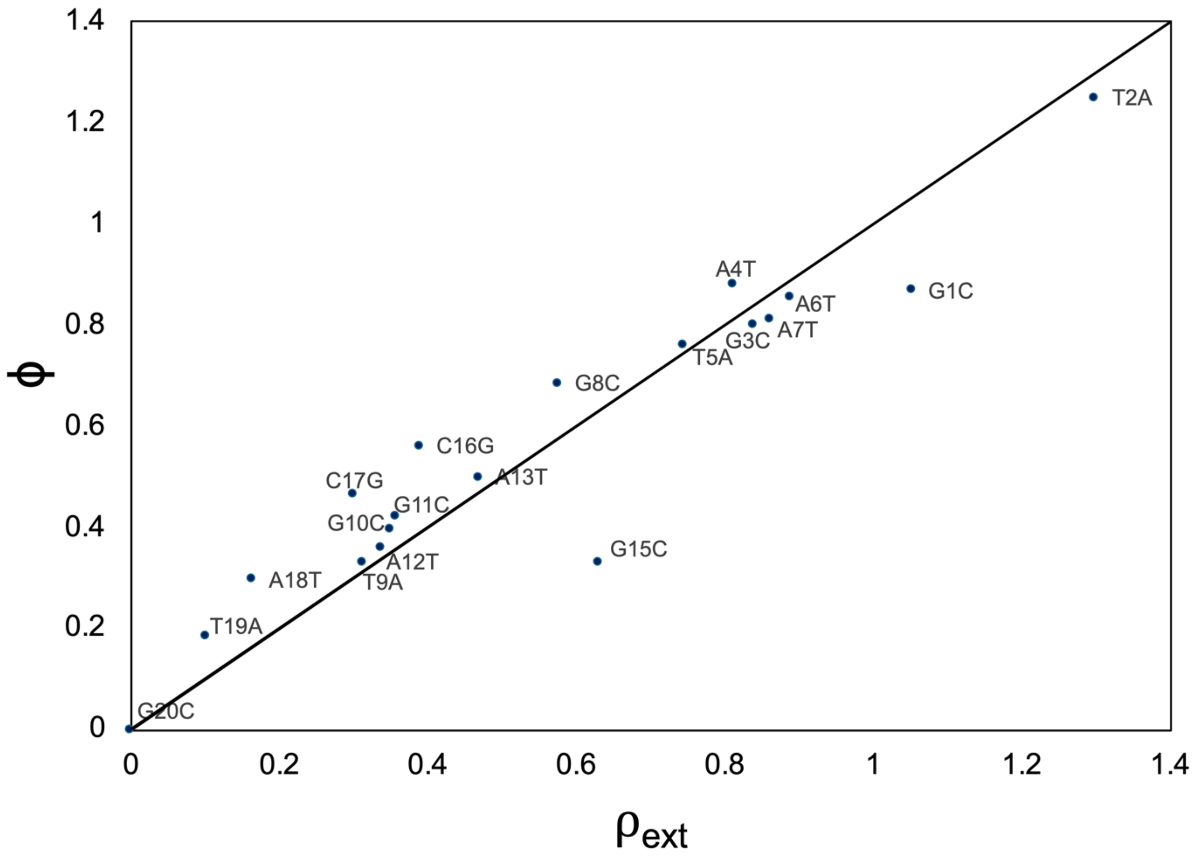


**Fig. S7.** Good general agreement between $\rho_{ext}$ values and $\phi$ values for Cas12a binding to DNA targets that include individual mismatches between the crRNA and the DNA target strand. The $\rho_{ext}$ value uses a value derived from nearest neighbor rules as a prediction for the thermodynamic effect of a given mismatch instead of an experimentally determined value, which for many RNA folding and assembly processes is be difficult to measure directly. Each mismatch in the plot is denoted by the nucleotide in the non-target strand. For example, the construct indicated as "T2A" has a sequence change at position 2, where T is changed to A on the non-target strand and A is changed to T on the target strand. The crRNA includes a U at position 2, generating a U-T mismatch between the crRNA and the DNA target strand. The line is the predicted 1:1 correspondence of the $\rho_{ext}$ values and $\phi$ values, and the data conform closely to the predictions in general, with an $R^2$ value of 0.86.

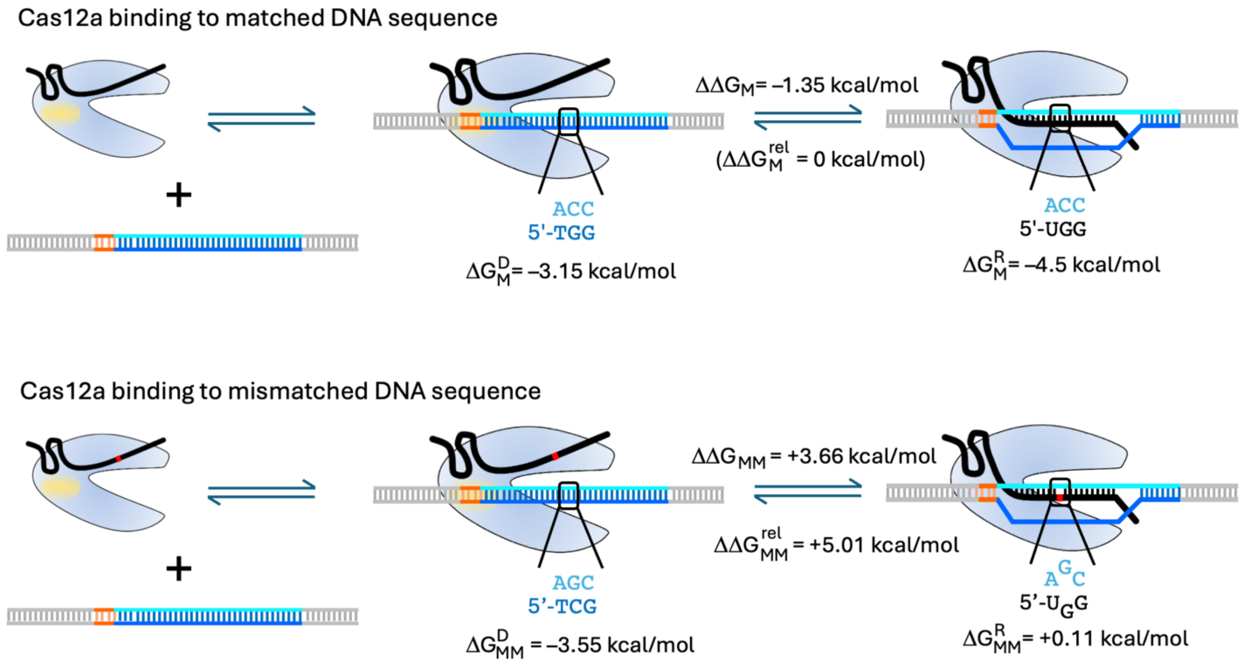


**Fig. S8.** Calculation of predicted free energy penalty for Cas12a binding to a DNA target that includes a mismatch between the crRNA and the DNA target strand. As an example, the sequences and numerical values for the prediction of the mismatch at position 10 are shown. Top, the two-step binding process for the matched DNA duplex. Bottom, the two-step binding process for the mismatched DNA duplex. In the first step, the Cas12a RNP associates with the DNA via the PAM contacts (shaded yellow on the protein, orange nucleotides in the DNA). In the second step, the R-loop is formed in a process that requires unwinding of the DNA. To calculate the predicted energetic penalty resulting from the mismatch, the predicted free energy change is calculated for each transition between the PAM-bound intermediate and the final complex. Only the nucleotide position being changed and its immediate neighbors are considered, as all the other positions are the same in the matched and mismatched duplexes. The predicted free energy change for formation each R-loop is calculated by subtracting the predicted ΔG for the DNA duplex from the predicted ΔG for the R-loop. To calculate the penalty, the predicted ΔG for R-loop formation with the matched DNA duplex (top) is subtracted from the predicted ΔG for R-loop formation with the mismatched DNA duplex (bottom) using the equation below:

$$\Delta\Delta G^{rel}_{MM} = \Delta G^{R}_{MM} - \Delta G^{D}_{MM} - (\Delta G^{R}_{M} - \Delta G^{D}_{M})$$

For the example sequence change shown:

$\Delta\Delta G^{rel}_{MM}$ = 0.11 kcal/mol – (–3.55 kcal/mol) – (–4.50 kcal/mol – (–3.15 kcal/mol))

$\Delta\Delta G^{rel}_{MM}$ = 0.11 kcal/mol + 3.55 kcal/mol + 4.50 kcal/mol – 3.15 kcal/mol

$\Delta\Delta G^{rel}_{MM}$ = 5.01 kcal/mol
